# Notch-Directed Germ Cell Proliferation Is Mediated by Proteoglycan-Dependent Transcription

**DOI:** 10.1101/2020.07.30.229997

**Authors:** Sandeep Gopal, Aqilah Amran, Andre Elton, Leelee Ng, Roger Pocock

## Abstract

Notch receptors are essential membrane-bound regulators of cell proliferation and differentiation in metazoa. In the nematode *Caenorhabditis elegans*, correct expression of GLP-1 (germline proliferation-1), a germline-expressed Notch receptor, is important for germ cell maintenance. However, mechanisms that regulate GLP-1 expression are undefined. Here, we demonstrate that an AP-2 transcription factor (APTF-2) regulates GLP-1 expression through calcium-dependent binding to a conserved motif in the *glp-1* promoter. Our data reveals that SDN-1 (syndecan-1), a transmembrane proteoglycan, regulates a TRP calcium channel in the soma to modulate the interaction between APTF-2 and *glp-1* promoter - thus providing a potential communication nexus between the germline and its somatic environment to control germ cell fate decisions.

## INTRODUCTION

Syndecans are a family of transmembrane proteoglycans that play important roles in metazoan development, regeneration, tissue repair and homeostasis (Afratis et al., 2017). Syndecans consist of conserved cytoplasmic and transmembrane domains and more divergent ectodomains that carry differentially sulfated glycosaminoglycan (GAG) chains (Choi et al., 2011). These GAG chains consist of heparan sulfate (HS) and chondroitin sulfate (CS) that can interact with ligands, extracellular matrix proteins, cytokines, growth factors and small molecules to coordinate signaling, cell adhesion and host defence mechanisms (Choi et al., 2011). Previously, we showed that syndecans control intrinsic cell behavior by regulating transient receptor potential canonical (TRPC) Ca^2+^ channels to maintain optimal Ca^2+^ influx over the plasma membrane (Gopal et al., 2015). Mechanistically, syndecans induce protein kinase C-alpha (PKCα)-dependent phosphorylation of a conserved serine residue in the cytoplasmic domain of TRPC channels (Gopal et al., 2015). This phosphorylation event closes the TRPC channel and thereby limits Ca^2+^ influx (Gopal et al., 2015). As Ca^2+^ can act as a secondary messenger, the control of Ca^2+^ influx by syndecans may enable communication between the environment, extracellular matrix and cytoplasmic/nuclear factors.

*Caenorhabditis elegans* encodes a single syndecan (SDN-1) that controls neuronal development, cell migration, behavior and spindle orientation (Dejima et al., 2014; Rhiner et al., 2005; Schwabiuk et al., 2009). Neuronal defects caused by loss of SDN-1 are suppressed by the removal of TRPC channels, revealing that Ca^2+^ regulation is an evolutionarily conserved function for syndecans (Gopal et al., 2015). Early studies of a *C. elegans sdn-1* loss-of-function mutant also revealed a reduction in brood size, suggesting a role for SDN-1 in progeny generation and germline function (Rhiner et al., 2005). The adult *C. elegans* hermaphrodite generates approximately 300 offspring from two syncytial germline tubes, within which germ cell nuclei are partially enclosed by membranes (Hubbard and Schedl, 2019). The germline is organised in a distal-to-proximal manner, with mitotically-dividing cells (including self-renewing germline stem cells - GSCs) within the distal progenitor zone (PZ), and proximal differentiated gametes (Hubbard and Schedl, 2019). As cells move through the germline from distal to proximal, they are influenced by distinct genetic programs that control their fate.

One of the most critical genetic regulators within the germline is the GLP-1 Notch receptor, which is required for establishing and maintaining the stem cell pool within the PZ (Austin and Kimble, 1987; Hubbard and Schedl, 2019; Kershner et al., 2013). In this context, a single-celled mesenchymal stem cell niche, called the distal tip cell (DTC), expresses the Notch ligands LAG-2 and APX-1 that trigger GLP-1 cleavage (Henderson et al., 1994; Nadarajan et al., 2009; Tax et al., 1994). The GLP-1 Notch intracellular domain that is released by this cleavage transcriptionally regulates downstream targets, including the redundantly-acting genes *lst-1* (lateral signaling target) and *sygl-1* (synthetic Glp), to maintain a pool of GSCs within the PZ (Kershner et al., 2014). Accurate regulation of GLP-1 expression and processing is critical for germline development: reduced GLP-1 expression causes germ cells to exit the mitotic cell cycle and inappropriately enter meiosis, whereas elevated GLP-1 signaling results in a germline tumor that is replete with mitotic cells (Austin and Kimble, 1987; Berry et al., 1997). Although much attention has been focused on deciphering mechanisms regulating Notch-ligand interactions and Notch processing, how Notch receptor transcription is controlled is not understood.

In this study, we reveal a critical function for SDN-1 in regulating germ cell fate by controlling GLP-1 expression. We found that SDN-1 acts non-cell autonomously from somatic gonadal sheath cells to promote GLP-1-dependent germ cell fate. Further, we reveal an important role for Ca^2+^ regulation in germ cell development as removal of the TRPC Ca^2+^ channel, TRP-2, suppresses germ cell fate defects caused by loss of SDN-1. Genetically downstream of SDN-1, we found that *glp-1* transcription is controlled through a deeply conserved promoter motif directly bound and regulated by the APTF-2 transcription factor. Binding of APTF-2 to the *glp-1* promoter is abrogated by elevated Ca^2+^, thus providing a mechanistic link between the SDN-1/TRP-2 axis and GLP-1 transcriptional regulation. Together, our data reveal a mechanism for soma-germline communication to control germ cell fate, where SDN-1 acts from somatic cells to control GLP-1 expression and germ cell behavior. This mode of regulation may enable interpretation and communication of extra-germline environmental cues, and thus optimize germ cell proliferation and progeny generation in ephemeral habitats and in pathophysiological conditions.

## RESULTS

### SDN-1 Controls Germ Cell Proliferation

In *C. elegans*, loss-of-function mutations in the sole syndecan (SDN-1) result in reduced progeny, suggesting aberrant germline function (Figure S1A) (Rhiner et al., 2005). To examine the *in vivo* importance of SDN-1 in the germline, we quantified germ cell number using three-dimensional germline reconstruction (Gopal et al., 2017). Using two independently-derived *sdn-1* deletion alleles, we found that loss of SDN-1 reduces the number of mitotically-dividing germ cells within the PZ, when compared to age-synchronized wild-type animals (Figure 1A-B). Furthermore, PZ length was shorter in *sdn-1* mutants (~16-18 cell diameters) compared to wild-type animals (~20 cell diameters) (Figure 1A and S1B). As SDN-1 functions during development to control multiple cell migration events, including DTC migration (Cram et al., 2006; Rhiner et al., 2005), we investigated whether post-developmental knockdown of *sdn-1* also causes a reduction in PZ germ cell number. We found that RNAi-mediated interference (RNAi) knockdown of *sdn-1* for 16 hours from the L4 larval stage (after DTC migration is complete) caused a reduction in PZ cell number (Figure S1C). These data support a post-developmental function for SDN-1 in controlling germ cell number. To assess the proliferative state of PZ cells in *sdn-1(zh20)* animals, we detected cells in M phase by staining germlines with an antibody against phospho-histone H3 (Figure 1C). We found that the number of M phase cells in *sdn-1(sh20)* animals were reduced by ~50% compared to wild-type (Figure 1C-D). These data reveal an important function for SDN-1 in regulating distal germ cell behavior.

**Figure 1.**
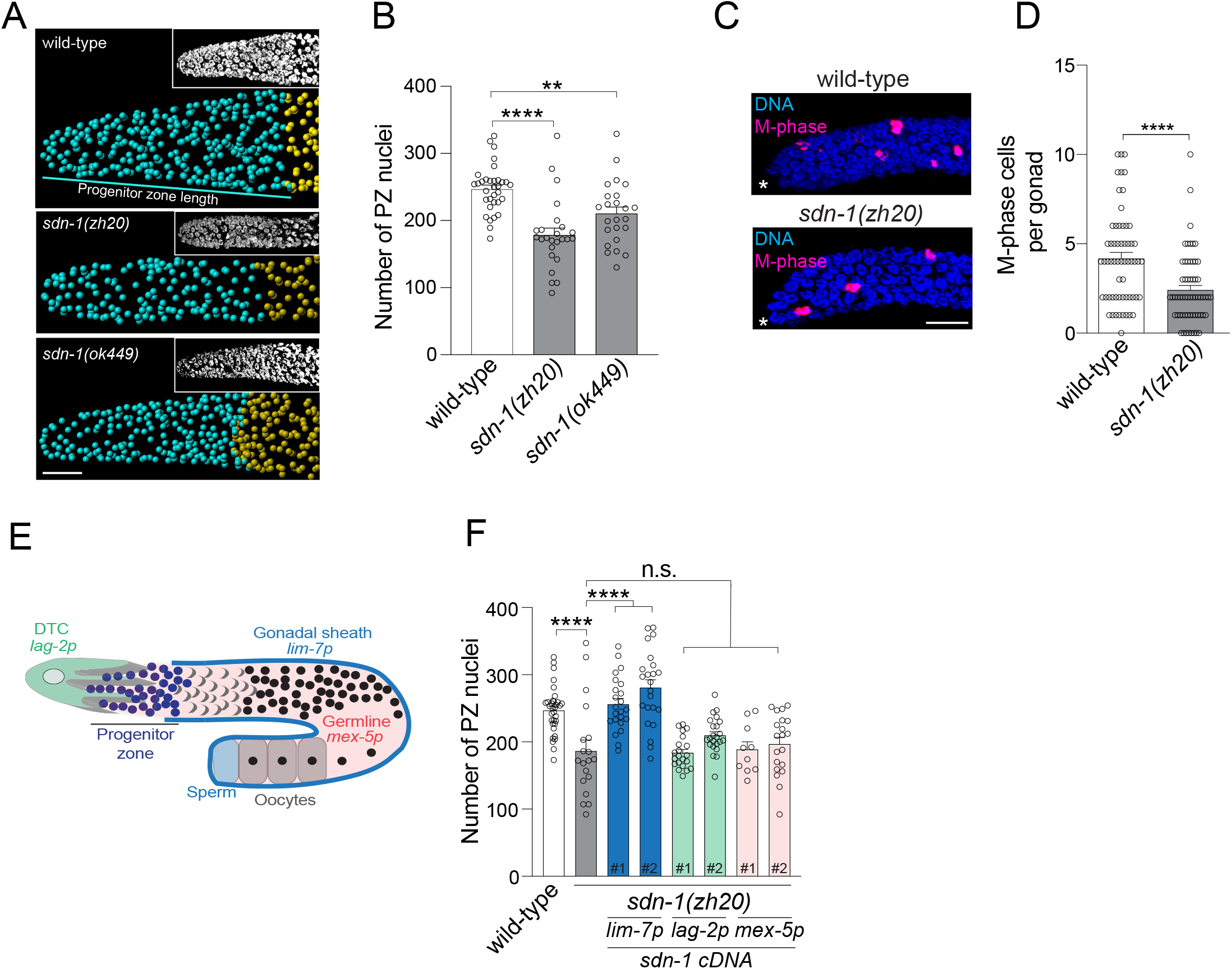
SDN-1 Controls Germ Cell Proliferation from the Somatic Sheath. (A-B) 3D germline analysis images (A) and quantification (B) of germline progenitor zone (PZ) nuclei (cyan spheres) in wild-type, *sdn-1(zh20)* and *sdn-1(ok449)* adult hermaphrodites. Insets in (A) = DAPI staining. Data are expressed as mean ± s.e.m. and statistical significance was assessed by ordinary one-way ANOVA. n>25. ****p<0.0001, **p<0.005. Scale bar = 10μm. (C-D) Immunostaining images (C) and quantification (D) of M-phase cells in distal germlines in wild-type and *sdn-1(zh20)* adult hermaphrodites. Germlines stained with DAPI to visualize DNA (blue) and anti-phospho-histone H3 (pH3) to visualize M-phase chromsomes (magenta). Data are expressed as mean ± s.e.m. and statistical significance was assessed by Welch’s t-test n=60. ****p<0.0001. Scale bar = 10μm. Asterisks, distal end of the gonad. (E-F) Schematic of one *C. elegans* germline arm (E) showing expression domains of promoters used in *sdn-1(zh20)* rescue experiments. Quantification of PZ nuclei (F) in wild-type, *sdn-1(zh20)* adult hermaphrodites +/− single-copy transgenic expression of *sdn-1* cDNA under tissue-specific promoters (*lag-2p* - DTC, *mex-5p* - germline, *lim-7p* gonadal sheath). Independent transgenic expression lines for each tissue-specific transgene is represented by #. Data are expressed as mean ± s.e.m. and statistical significance was assessed by ordinary one-way ANOVA. n>20. ****p<0.0001, n.s. = not significant.

Expression of *sdn-1* in the germline or associated somatic tissues has not been detected with transgenic reporters (Minniti et al., 2004; Rhiner et al., 2005; Schwabiuk et al., 2009). However, spatiotemporal sequencing analysis shows that *sdn-1* mRNA is detected at low levels in hermaphrodite germline tissue (Diag et al., 2018; Tzur et al., 2018). To characterize where SDN-1 acts to control germ cell behavior, we used defined tissue-specific promoters to express a single copy of *sdn-1* cDNA, using miniMos-mediated genomic insertion (Figure 1E-F) (Frokjaer-Jensen et al., 2014). We expressed *sdn-1* cDNA in either the germline (*mex-5* promoter), DTC (*lag-2* promoter) or gonadal sheath (*lim-7* promoter) of *sdn-1(zh20)* mutant animals and quantified PZ cell number (Figure 1E-F) (Hall et al., 1999; Henderson et al., 1994; Merritt et al., 2008). We obtained two independent integrated lines for each single-copy transgene and found that expressing *sdn-1* only in the gonadal sheath rescued the *sdn-1(zh20)* PZ germ cell number phenotype (Figure 1F). Similarly, expression of *sdn-1* in the gonadal sheath rescued the number of M phase cells in *sdn-1(sh20)* animals to wild-type levels (Figure S2A). These data show that SDN-1 acts non-cell autonomously from the somatic gonad sheath cells to control germ cell proliferation.

The gonadal sheath consists of somatic cells that cover the surface of the germline meiotic region and also send processes into the proximal part of the PZ (Hall et al., 1999; Killian and Hubbard, 2005; McCarter et al., 1997). Our finding that SDN-1 functions in gonadal sheath cells to control germ cell behavior suggests that SDN-1 may interpret extra-germline signals (e.g. environmental and matrix cues) at the germline-soma interface. Syndecans have established roles in regulating metazoan cell behavior through binding to extracellular ligands via GAG chains that are attached to three conserved serine residues in their ectodomain (Esko and Zhang, 1996; Gopal et al., 2010). We therefore investigated whether GAG chain attachment is important for SDN-1 regulation of PZ germ cell number. We mutated all three SDN-1 GAG attachment sites (Minniti 2004) from serine to alanine (SDN-1-AAA) and expressed a single copy of SDN-1-AAA in the gonadal sheath of *sdn-1(zh20)* animals (Figure S2B-D). We found that unlike wild-type SDN-1, GAG chain-deficient SDN-1-AAA was unable to rescue the PZ germ cell length or M phase cell phenotype of *sdn-1(zh20)* animals (Figure S2C-D). Collectively, these data show that SDN-1 acts from the gonadal sheath to control PZ germ cell behavior and requires attachment of extracellular GAG chains to perform this function.

### SDN-1 Regulates Germ Cell Fate Through the GLP-1 Notch Receptor

The GLP-1 Notch receptor is a major regulator of germ cell fate in the *C. elegans* germline (Hubbard and Schedl, 2019). GLP-1 depletion causes germ cells to prematurely exit the mitotic cell cycle and as such reduces PZ cell number (Hubbard and Schedl, 2019). We therefore asked whether *sdn-1* acts in the same pathway as *glp-1* to control PZ cell number. We knocked down *glp-1* for 16 hours from the L4 stage using RNAi and analysed young adult hermaphrodites. As shown previously, decreased *glp-1* expression in wild-type animals reduces PZ cell number (Figure 2A) (Austin and Kimble, 1987). However, *glp-1* knockdown did not reduce PZ cell number of *sdn-1(zh20)* mutant animals suggesting that *sdn-1* and *glp-1* act in the same genetic pathway (Figure 2A). In support of this, we found that a *glp-1(e2141)* loss-of-function mutant also did not further reduce the number of PZ cells or M phase cells of *sdn-1(zh20)* mutant animals (Figure S3A-B). Together, these data suggest that *glp-1* and *sdn-1* act in the same genetic pathway to control PZ cell number.

**Figure 2.**
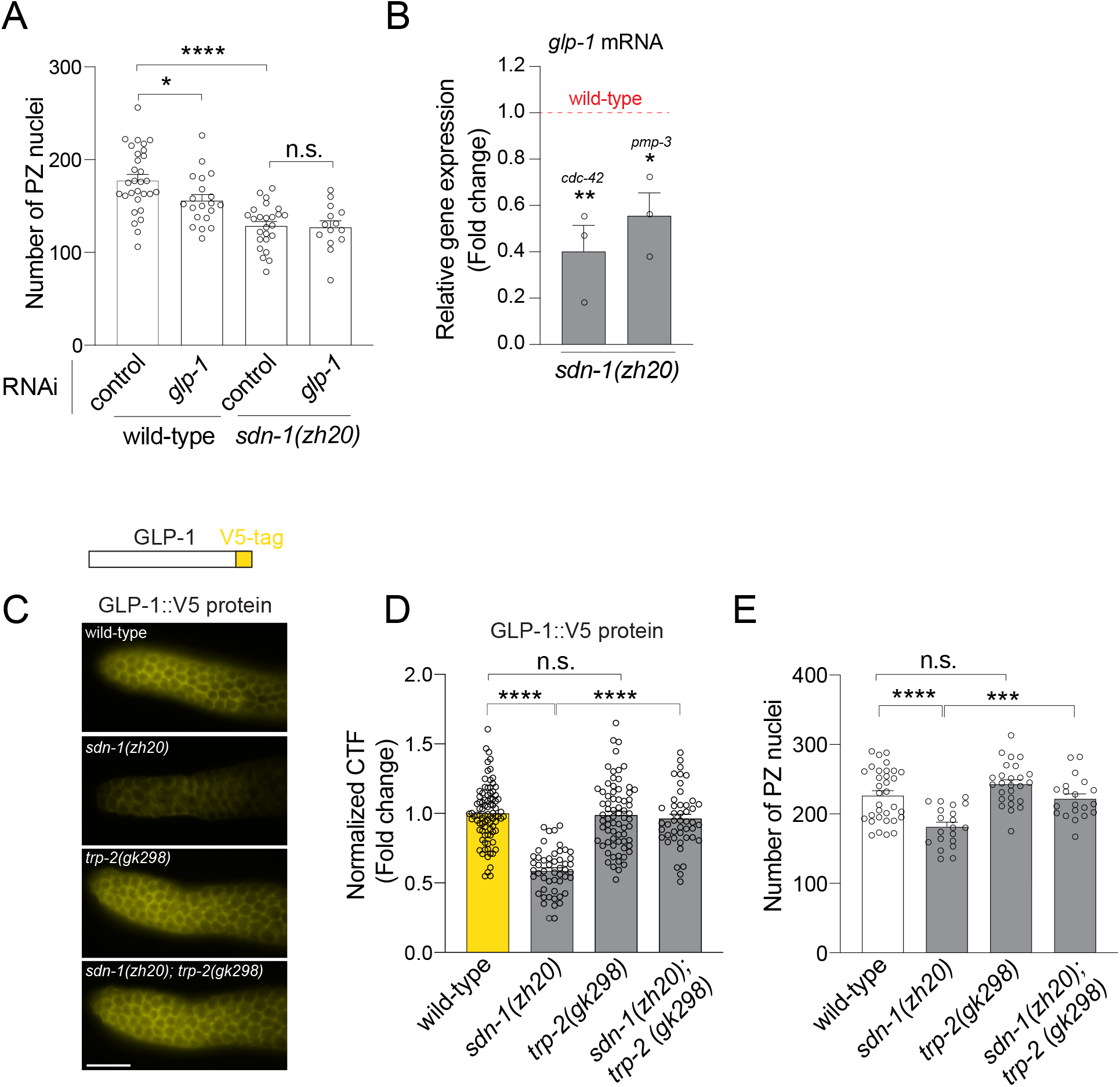
SDN-1 Promotes GLP-1/Notch Expression to Control Mitotic Germ Cell Fate. (A) Quantification of PZ nuclei in wild-type and *sdn-1(zh20)* adult hermaphrodites grown on control (L4440 vector) and *glp-1* RNAi bacteria. Data are expressed as mean ± s.e.m. and statistical significance was assessed by ordinary one-way ANOVA. n>15. ****p<0.0001, *p<0.05, n.s. = not significant. (B) Quantification of *glp-1* mRNA by qPCR in *sdn-1(zh20)* animals compared to wild-type. Two independent reference genes (*cdc-42*) and (*pmp-3*) were used to normalize expression. RNA was isolated fom dissected germlines. Data are expressed as mean ± s.e.m. and statistical significance was assessed by ordinary one-way ANOVA. n=3 independent biological replicate RNA RNA samples for each genotype. **p<0.01, *p<0.05. (C-D) Schematic of GLP-1::V5 protein and immunofluorescence images (C) and quantification (D) of GLP-1::V5 expression in the PZ of wild-type, *sdn-1(zh20)*, *trp-2(gk298)* and *sdn-1(zh20); trp-2(gk298)* adult hermaphrodites. Data are expressed as mean ± s.e.m. and statistical significance was assessed by ordinary one-way ANOVA. n>45. ****p<0.0001, n.s. = not significant. Scale bar = 10μm (E) Quantification of PZ nuclei in wild-type, *sdn-1(zh20)*, *trp-2(gk298)* and *sdn-1(zh20); trp-2(gk298)* adult hermaphrodites. Data are expressed as mean ± s.e.m. and statistical significance was assessed by ordinary one-way ANOVA. n>20. ****p<0.0001, ***p<0.001, n.s. = not significant.

To investigate the potential mechanistic relationship between *sdn-1* and *glp-1*, we measured *glp-1* mRNA and protein in *sdn-1(zh20)* mutant animals. First, we measured *glp-1* mRNA from extruded germlines of wild-type and *sdn-1(zh20)* animals using quantitative real-time PCR (qPCR). We found that *glp-1* mRNA levels are reduced by ~50% in *sdn-1(zh20)* mutant animals (Figure 2B). To enable quantification of endogenous GLP-1 protein, we utilized a CRISPR-Cas9 generated strain in which GLP-1 is endogenously tagged with the viral V5 epitope (GLP-1::V5) (Figure 2C) (Sorensen et al., 2020). The V5 tag does not detrimentally affect germline function, with GLP-1::V5 hermaphrodites having a similar number of PZ germ cells to wild-type (Figure S4A). Immunofluorescence detection of GLP-1::V5, using an anti-V5 antibody, showed prominent expression at the distal end of the germline and in embryos, as shown previously with anti-GLP-1 immunofluorescence (Figure 2C and S4B) (Crittenden et al., 1994; Sorensen et al., 2020). We quantified GLP-1::V5 expression in the distal end of the germline and found that loss of *sdn-1* reduced the expression of GLP-1::V5 protein by ~50% (Figure 2C-D). Together, these data suggest that SDN-1 regulates PZ germ cell number by signaling from the somatic sheath to positively regulate *glp-1* expression in the germline.

Next, we investigated how SDN-1-mediated signaling and GLP-1 expression are linked. Our previous studies using vertebrate and invertebrate models showed that the canonical role of syndecans in controlling cell behavior is to repress transient receptor potential canonical (TRPC) channel function, and thereby limit cellular Ca^2+^ influx (Gopal et al., 2015). We therefore investigated whether SDN-1 uses this mechanism to control *glp-1* expression. Three TRPC-like channels are encoded by *C. elegans*, of which only TRP-2 is expressed robustly in germline-associated tissue outside the gametes (Tzur et al., 2018). Based on our previous study (Gopal et al., 2015), we posited that introduction of the *trp-2(gk298)* mutation into *sdn-1(zh20)* animals would restore GLP-1::V5 expression to wild-type levels. Indeed, we found that the reduction of GLP-1::V5 expression in *sdn-1(zh20)* animals is restored to wild-type levels when *trp-2* is deleted (Figure 2C-D). In addition, the reduced PZ germ cell number phenotype of *sdn-1(zh20)* animals is suppressed in *sdn-1(zh20); trp-2(gk298)* double mutant animals (Figure 2E). Together, these data reveal that the SDN-1/TRP-2 regulatory axis, acting from the gonadal sheath, positively regulates *glp-1* expression to influence PZ germ cell proliferation.

### *glp-1* Transcription is Regulated Through a Conserved Promoter Motif

How does SDN-1 signaling control *glp-1* expression? We reasoned that SDN-1 regulates *glp-1* expression at the level of transcription because the reduction of *glp-1* mRNA mirrors the reduction of GLP-1 protein in *sdn-1(zh20)* mutant animals (Figure 2). However, transcriptional mechanisms controlling Notch receptor are unknown.

To elucidate how *glp-1* is transcriptionally regulated, we surveyed *glp-1* promoters from four *Caenorhabditis* species for conserved DNA sequences and identified a highly conserved 9 base pair (bp) sequence (Figure 3A). In the *C. elegans* promoter, the conserved sequence (TGCCACCCG) is located 140 bp upstream of the ATG start codon, suggesting that this motif may be a binding site for transcriptional regulators. Bioinformatic analysis revealed that this conserved *glp-1* promoter motif contains a consensus sequence for the transcription factor AP-2 (TFAP2) family. To interrogate the importance of this conserved motif for GLP-1 expression and function we used the endogenously tagged GLP-1::V5 strain. Using CRISPR-Cas9 editing, we introduced a 5 bp deletion within the conserved sequence (*glp-1promΔ*) and examined GLP-1::V5 expression and PZ germ cell number. Using anti-V5 immunofluorescence, we found that deleting the putative TFAP2 motif reduced GLP-1::V5 levels in the germline PZ (Figure 3B-C). To confirm the importance of the putative TFAP2 motif for *glp-1::v5* mRNA expression we used single-molecule fluorescence *in situ* hybridization (smFISH) to detect *glp-1::V5* transcripts. We found that *glp-1::v5* transcript levels in the germline are too low for comparison due to background fluorescence, however, *glp-1::v5* is robustly detected in 2-cell embryos. We found that deletion of the putative TFAP2 motif reduced *glp-1* transcript levels in 2-cell embryos, revealing that this motif is required for correct *glp-1* expression in both embryos and the germline. (Figure S5A-B).

**Figure 3.**
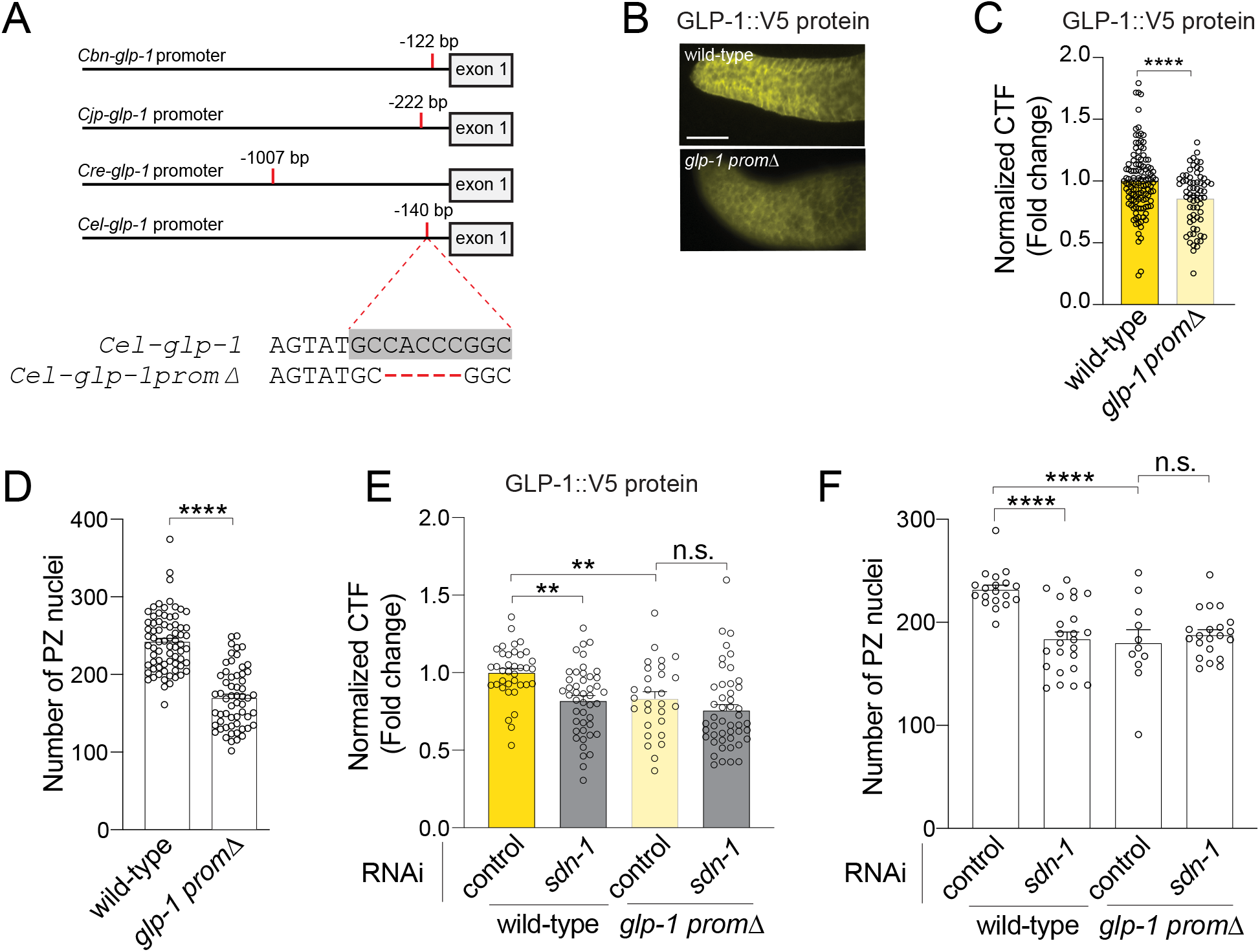
A Conserved Promoter Motif Controls GLP-1 Expression and Germ Cell Number. (A) Location of a conserved motif in the *glp-1* promoter of four *Caenorhabditis* species. *Cbn* = *C. brenneri, Cjp* = *C. japonica*, *Cre* = *C. remanei*, *Cel* = *C. elegans*. The conserved motif in *C. elegans glp-1* promoter is highlighted in grey. Deleted bases in CRISPR-engineered *glp-1* promoter mutant in red. (B-C) Immunofluorescence images (B) and quantification (C) of GLP-1::V5 protein in the distal germline of wild-type and *glp-1prom*Δ (5 bp deletion in conserved promoter motif) adult hermaphrodites. Data are expressed as mean ± s.e.m. and statistical significance was assessed by unpaired t-test. n>60. ****p<0.0001. Scale bar = 10μm (D) Quantification of PZ nuclei of wild-type and *glp-1prom*Δ adult hermaphrodites. Data are expressed as mean ± s.e.m. and statistical significance was assessed by unpaired t-test. n>45. ****p<0.0001. (E) Quantification of GLP-1::V5 protein in the distal germline of adult hermaphrodites (+/− *sdn-1* RNAi) expressing wild-type *glp-1* or *glp-1prom*Δ adult hermaphrodites. Data are expressed as mean ± s.e.m. and statistical significance was assessed by ordinary one-way ANOVA. n>25. **p<0.01, n.s. = not significant. (F) Quantification of PZ nuclei in the distal germline of adult hermaphrodites (+/− *sdn-1* RNAi) expressing wild-type *glp-1* or *glp-1prom*Δ. Data are expressed as mean ± s.e.m. and statistical significance was assessed by ordinary one-way ANOVA. n>10. ****p<0.0001, n.s. = not significant.

Next, we investigated the functional importance of the putative TFAP2 motif by quantifying PZ germ cell number in wild-type *glp-1::v5* and mutant *glp-1promΔ* animals (Figure 3D). We found that deletion of the putative TFAP2 motif reduced the number of PZ germ cells by ~30% (Figure 3D). Our data suggest that SDN-1 signaling controls *glp-1* transcription, therefore we examined whether SDN-1 may act through the putative TFAP2 motif. If SDN-1 controls *glp-1* expression through the putative TFAP2 motif, we would not expect to observe an additive effect when the *glp-1promΔ* mutant was combined with the *sdn-1(zh20)* mutant strain. In congruence, *glp-1promΔ* and *glp-1promΔ; sdn-1* RNAi animals exhibit equivalent reduction in GLP-1::V5 expression and PZ cell number (Figure 3E-F). Whereas, *sdn-1* RNAi reduces GLP-1::V5 expression and PZ cell number in animals that harbor a wild-type *glp-1* promoter (Figure 3E-F). Together, these data identify a highly conserved putative TFAP2 motif within the *glp-1* promoter that is important for regulation of *glp-1* expression and germ cell fate.

We have shown that deletion of putative TFAP2 motif reduces GLP-1 expression and PZ germ cell number (Figure 3). Next, we used CRISPR-Cas9 to mutate two bases in the putative TFAP2 binding motif of the *glp-1::v5* promoter (*glp-1prom(aa)*) that are predicted to disrupt TFAP2 binding (Figure 4A) (Woodfield et al., 2010). Animals harboring these two base pair mutations in the *glp-1* promoter exhibited reduced GLP-1::V5 expression and PZ germ cell number to a similar extent as the motif deletion (Figure S5C-D), supporting a potential role for TFAP2-mediated regulation of *glp-1* expression.

**Figure 4.**
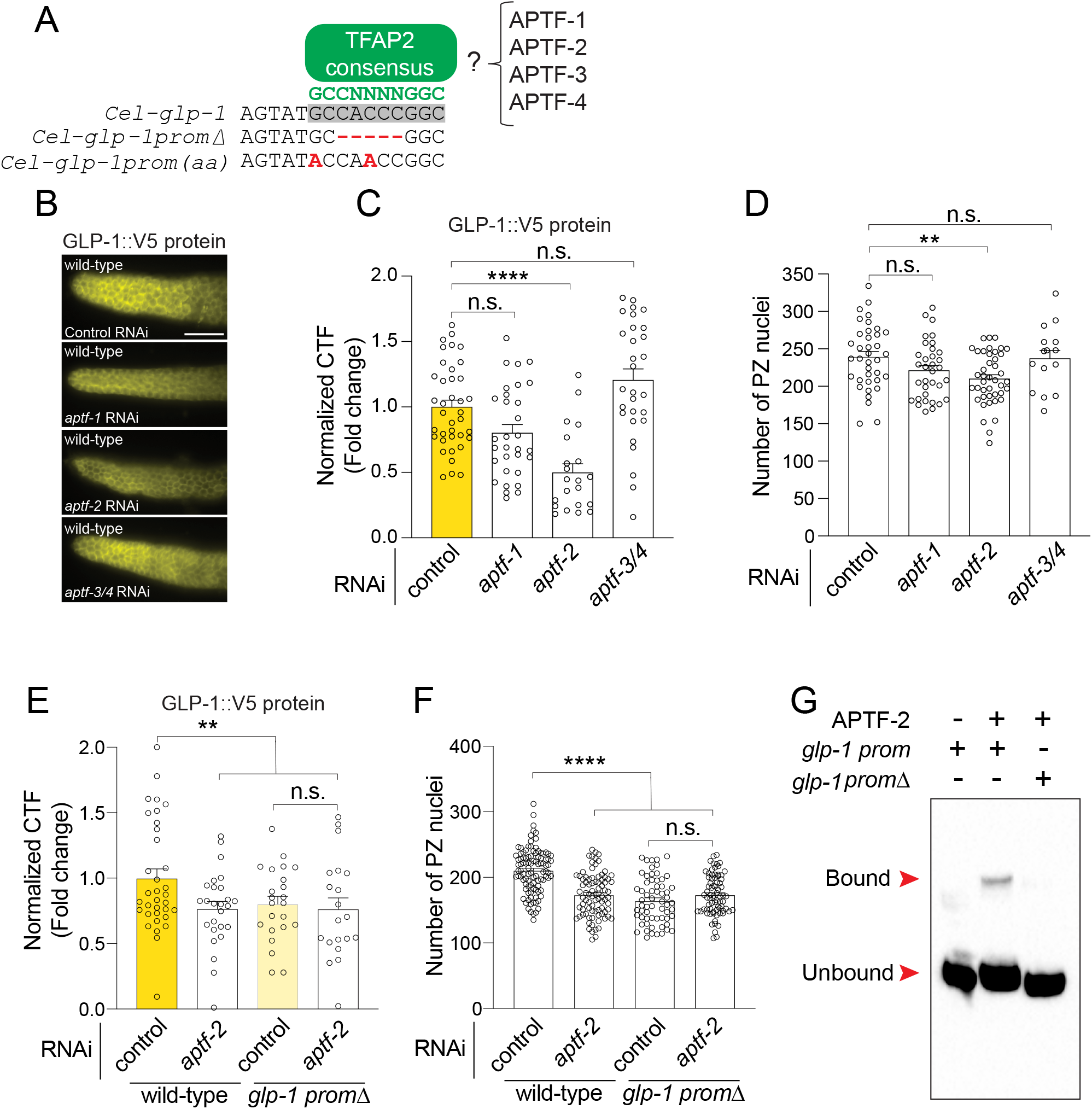
*glp-1* Transcription is Directly Regulated by APTF-2. (A) A consensus binding site (GCCNNNNGGC) for AP-2 transcription factors is located within the conserved *glp-1* promoter motif. TFAP2 consensus motif (green), TFAP2 motif in the *glp-1* promoter (grey box), *glp-1* promoter deletion/mutations (red). (B-C) Immunofluorescence images (B) and quantification (C) of GLP-1::V5 protein in the distal germline of wild-type adult hermaphrodites following RNAi knockdown of *aptf-1*, *aptf-2* and *aptf-3/4*. Data are expressed as mean ± s.e.m. and statistical significance was assessed by ordinary one-way ANOVA. n>20. ****p<0.0001, n.s. = not significant. Scale bar = 10μm. (D) Quantification of PZ nuclei in wild-type adult hermaphrodites following RNAi knockdown of *aptf-1*, *aptf-2* and *aptf-3/4*. Data are expressed as mean ± s.e.m. and statistical significance was assessed by ordinary one-way ANOVA. n>15. **p<0.01, n.s. = not significant. (E) Quantification of GLP-1::V5 protein in the distal germline of adult hermaphrodites (+/− *aptf-2* RNAi) expressing wild-type *glp-1* or *glp-1prom*Δ. Data are expressed as mean ± s.e.m. and statistical significance was assessed by ordinary one-way ANOVA. n>20. **p<0.01, n.s. = not significant. (F) Quantification of PZ nuclei in the distal germline of adult hermaphrodites (+/− *aptf-2* RNAi) expressing wild-type *glp-1* or *glp-1prom*Δ. Data are expressed as mean ± s.e.m. and statistical significance was assessed by ordinary one-way ANOVA. n>30. ****p<0.0001, n.s. = not significant. Scale bar = 10μm (G) EMSA of APTF-2::V5 protein produced in HEK293T cells and the TFAP2 consensus motif within the *glp-1* promoter. DNA bound and unbound with APTF-2::V5 is marked. *glp-1* prom = wild-type and *glp-1* promΔ = TFAP2 motif deletion.

### APTF-2 Directly Regulates *glp-1* Expression

*C. elegans* encodes four TFAP2 family members (APTF-1-4), of which APTF-2 and APTF-3 expression has been detected in the germline (Diag et al., 2018; Tzur et al., 2018). To determine the importance of APTF-1-4 in regulating *glp-1* expression, we used RNAi to individually knock down their expression in wild-type animals for 16 hours from the L4 larval stage and measured GLP-1::V5 levels in the adult germline (Figure 4B-C). We found that *aptf-2* RNAi reduced GLP-1::V5 expression in the distal end of the germline (Figure 4B-C). In addition, *aptf-2* RNAi lowered PZ germ cell number, as did an *aptf-2(qm27)* mutant that harbors a point mutation in the APTF-2 DNA binding domain (Figure 4D and S6) (Budirahardja et al., 2016). In contrast, RNAi knockdown of *aptf-1* and *aptf-3/4* caused non-significant changes to GLP-1::V5 expression and PZ germ cell number (Figure 4B-D and S6).

We found that *aptf-2* RNAi knockdown did not further reduce GLP-1::V5 expression or PZ germ cell number in the germline of animals in which the TFAP2 binding site in the *glp-1* promoter is deleted (Figure 4E-F). This suggests that APTF-2 controls *glp-1* expression and function by directly interacting with the conserved TFAP2 binding site. We tested this assertion using electrophoretic mobility shift assays (EMSAs). First, we expressed and purified APTF-2::V5-containing nuclear fractions from mammalian HEK293T cells. Next, we asked whether APTF-2::V5 could cause a shift in biotin-labelled *glp-1* promoter migration (Figure 4G). We found that APTF-2::V5 indeed binds to the functionally-important TFAP2 motif within the *glp-1* promoter and that this interaction is competed away with unlabeled *glp-1* promoter (Figure 4G and S7). Further, deletion of the TFAP2 motif abrogates APTF-2 binding to the *glp-1* promoter, confirming its importance in regulating *glp-*1 expression (Figure 4G). Taken together, these data show that APTF-2 directly binds to the *glp-1* promoter to drive its expression and function in the *C. elegans* germline.

### APTF-2 Regulation of GLP-1 Requires SDN-1

Our data show that APTF-2 positively regulates *glp-1* expression by directly binding to the *glp-1* promoter. Previous *in vitro* studies in mammalian cells revealed that the transcriptional activity of APTFs can be modulated by Ca^2+^ (Deyama et al., 1999). We therefore posited that regulation of Ca^2+^ levels by the SDN-1/TRP-2 axis may control the ability of APTF-2 to regulate *glp-1* expression in the *C. elegans* germline. To examine this possibility, we performed genetic analysis to determine whether regulation of GLP-1::V5 expression and PZ germ cell number by APTF-2 is dependent on SDN-1/TRP-2.

If SDN-1 controls PZ germ cell number through the APTF-2 regulation of GLP-1, we would expect that SDN-1 regulation of *glp-1* expression is APTF-2-dependent. To examine this, we measured GLP-1::V5 levels in wild-type and *sdn-1(zh20)* animals after RNAi knockdown of *aptf-2.* We found that GLP-1::V5 levels are reduced in *sdn-1(zh20)* animals, however, this was not further reduced by *aptf-2* RNAi knockdown (Figure 5A). To examine the biological relevance of this finding, we performed *aptf-2* RNAi knockdown in *sdn-1(zh20)* mutant animals and analyzed PZ germ cell number. We found that unlike in wild-type animals, no reduction in PZ germ cell number was observed in *sdn-1(zh20)* mutant animals following RNAi knockdown of *aptf-2* (Figure 5B). These data suggest that *sdn-1* acts in the same genetic pathway as *aptf-2* and *glp-1* to control PZ germ cell number.

**Figure 5.**
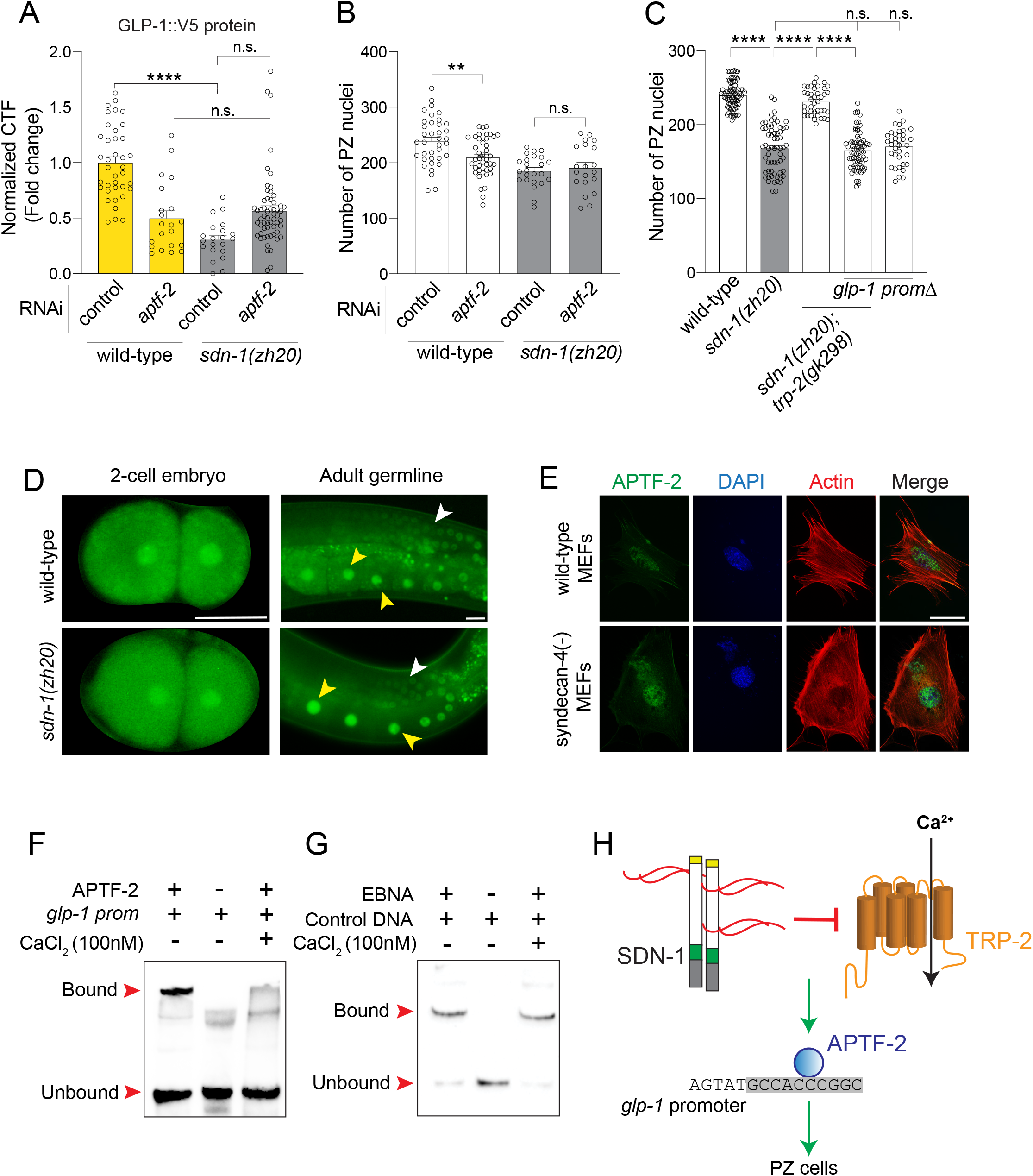
Calcium Controls APTF-2 Binding to the *glp-1* Promoter. (A) Quantification of GLP-1::V5 protein in the distal germline of wild-type and *sdn-1(zh20)* adult hermaphrodites (+/− *aptf-2* RNAi). Data are expressed as mean ± s.e.m. and statistical significance was assessed by ordinary one-way ANOVA. n>20. ****p<0.0001, n.s. = not significant. (B) Quantification of PZ nuclei in wild-type and *sdn-1(zh20)* adult hermaphrodites (+/− *aptf-2* RNAi). Data are expressed as mean ± s.e.m. and statistical significance was assessed by ordinary one-way ANOVA. n>20. **p<0.01, n.s. = not significant. (C) Quantification of PZ nuclei in wild-type, *sdn-1(zh20)* and *sdn-1(zh20); trp-2(gk298)* adult hermaphrodites expressing wild-type *glp-1* or *glp-1prom*Δ. n = >35. ****p<0.0001, n.s. = not significant. (D) Expression pattern of ATPF-2::GFP in 2-cell embryos and the adult germline of wild-type and *sdn-1(zh20)* hermaphrodites. APTF-2::GFP in the germline (white arrowheads) and oocytes (yellow arrowheads). Scale bar 25μm. (E) Immunofluorescence micrographs of wild-type and syndecan-4 knockout mouse embryonic fibroblasts (MEFs) expressing APTF-2::V5, co-stained with DAPI (DNA) and phalloidin (actin). APTF-2::V5 is nuclear-localized in both wild type and syndecan-4(-) cells, which show characteristically disorganized cytoskeleton. Scale bar = 50μm. (F) EMSA of APTF-2::V5 protein produced in HEK293T cells and the TFAP2 consensus motif within the *glp-1* promoter (+/− 100nM CaCl_2_). DNA bound and unbound with APTF-2::V5 is marked (red arrowheads). *glp-1* prom = wild-type. (G) EMSA of Epstein-Barr Nuclear Antigen Extract (EBNA) protein extract and the control DNA duplex (+/− 100nM CaCl_2_). DNA bound and unbound with EBNA is marked. (H) Mechanism for SDN-1 regulation of germ cell proliferation. Regulation of the TRP-2 calcium channel by SDN-1 controls the ability of APTF-2 binding to a conserved motif on the *glp-1* promoter.

We have shown that the reduction PZ germ cell number in *sdn-1(zh20)* animals is suppressed by removal of the TRP-2 channel (Figure 2E). If SDN-1 controls PZ germ cell number genetically upstream of APTF-2, deleting the APTF-2 binding site in the *glp-1* promoter would abrogate the suppressive effect of TRP-2 loss on the *sdn-1(zh20)* mutant phenotype. Indeed, when the APTF-2 binding site in the *glp-1* promoter is deleted, removal of TRP-2 does not suppress the *sdn-1(zh20)* mutant reduction of PZ germ cell number (Figure 5C). Together, these data posit that modulation of Ca^2+^ levels by the SDN-1/TRP-2 axis controls APTF-2-dependent regulation of PZ germ cell number by GLP-1.

### Calcium Regulates APTF-2 Binding to the *glp-1* Promoter

Mechanistically, changes in the level of cellular Ca^2+^ have been shown to affect the subcellular localization of transcription factors as well as their affinity for gene regulatory regions (Beals et al., 1997; Corneliussen et al., 1994; Mao and Wiedmann, 1999). We therefore examined these scenarios in the context of APTF-2 using *C. elegans in vivo* and mammalian cell-based methods.

To investigate whether SDN-1 regulates the expression of APTF-2, we used CRISPR-Cas9 to knock-in green fluorescent protein (*gfp)* coding sequence to the 3’ end of the *aptf-2* gene (Figure S8A). APTF-2::GFP reporter expression was confirmed by Western blot and fluorescence microscopy, revealing nuclear expression in the germline, oocytes and embryos (Figures 5D and S8B-D). We found however that the localization of endogenous APTF-2::GFP expression is unchanged in *sdn-1(zh20)* animals (Figure 5D). Next, using mouse embryonic fibroblasts that lack syndecan-4, and thus have constitutively high levels of intracellular Ca^2+^ due to dysregulation of TRPC7 channels (Gopal et al., 2015), we found that APTF-2::V5 nuclear localization is not overtly affected (Figure 5E). These data show that high intracellular Ca^2+^ does not overtly alter APTF-2 subcellular localization. Next, we used EMSA to investigate whether Ca^2+^ controls the affinity of APTF-2::V5 to the *glp-1* promoter. We found that binding of APTF-2::V5 to the *glp-1* promoter, but not a control protein-DNA interaction, was inhibited by an increase Ca^2+^ levels (Figure 5F-G). Whether Ca^2+^ levels directly affect the ability of APTF-2 to interact with the *glp-1* promoter or whether Ca^2+^ modulates an intermediate regulator of APTF-2 is yet to be discovered.

Our collective data reveal a mechanism where control of a TRP-2 channel by the transmembrane proteoglycan SDN-1 regulates GLP-1 expression and germ cell fate (Figure 5H). Within the germline, and genetically downstream of SDN-1/TRP-2, the APTF-2 transcription factor directly regulates *glp-1* expression through a conserved TFAP2 binding site. APTF-2 binding to the *glp-1* promoter is sensitive to Ca^2+^, providing a mechanistic link between the SDN-1/TRP-2 function in the gonadal sheath and *glp-1* germline transcription.

## DISCUSSION

Here we have identified a mechanism by which the transmembrane proteoglycan SDN-1 controls germ cell fate by regulating the expression of the GLP-1 Notch receptor. SDN-1 performs this function non-cell autonomously from somatic gonadal sheath cells that surround the germline. We found that the established role of SDN-1 in controlling transient receptor potential calcium channels is required for correct GLP-1 expression and germ cell fate. Further, SDN-1 requires the attachment of extracellular sugar chains to control germ cell fate, suggesting that extracellular ligands, environmental cues or matrix proteins control this SDN-1/TRP-2 axis. In the germline, GLP-1 expression is directly regulated by the APTF-2 transcription factor through a conserved DNA binding motif within the *glp-1* promoter, an interaction that is dependent on the level of cellular Ca^2+^. Therefore, our cumulative data reveal a mechanism of soma-germline signaling to promote mitotic germ cell fate by regulating GLP-1 expression.

Notch signaling is an ancient and highly conserved mechanism used in metazoa to control cell-cell communication. Notch receptor mutants were first discovered in *Drosophila* and named based on the dominant notched wing phenotype observed (Dexter, 1914). The advent of molecular genetics enabled fine dissection of the Notch signaling pathway and its importance in development and disease (Andersson et al., 2011; Greenwald, 1985; Greenwald et al., 1983; Kidd et al., 1986; Kopan and Ilagan, 2009; Wharton et al., 1985). For example, seminal discoveries revealing the function and processing of the GLP-1 and LIN-12 Notch receptors in *C. elegans* have contributed considerably to our understanding of this important signaling pathway (Austin and Kimble, 1987, 1989; Evans et al., 1994; Greenwald et al., 1983; Levitan and Greenwald, 1995; Shaye and Greenwald, 2002; Yochem and Greenwald, 1989). In the context of the *C. elegans* germline, it has been known for decades that GLP-1 signaling intensity is important for controlling mitotic cell fate (Austin and Kimble, 1987; Berry et al., 1997). However, elucidation of mechanisms controlling *glp-1* transcription has been lacking. Our study therefore identifies control of *glp-1* transcription as an additional layer of regulation to the Notch signaling pathway in the *C. elegans* germline and suggests similar regulatory mechanisms exist in other organisms.

Previous reports have implicated proteoglycans and GAG chain modifications in the regulation and differentiation of embryonic and adult stem cells in mammals (Johnson et al., 2007; Kraushaar et al., 2013; Nairn et al., 2007; Pisconti et al., 2010). During embryonic stem cell (ESC) differentiation, distinct sulfation patterns in surface GAGs are observed - from low sulfation in pluripotent cells to high sulfation in differentiated cells (Hirano et al., 2012; Johnson et al., 2007). Further, GAG chain-deficient ESCs are unable to differentiate, revealing a function for GAG dynamics in controlling stem cell differentiation (Johnson et al., 2007). In adult mammalian muscle, it was also shown that Syndecan-3 is required for satellite cell proliferation (Pisconti et al., 2010). Intriguingly, defective cellular behavior of Syndecan-3-null satellite cells is rescued by expression of a constitutively active Notch intracellular domain (Pisconti et al., 2010). However, in this context, Syndecan-3 directly interacts with the Notch receptor and is required for Notch processing (Pisconti et al., 2010). GAGs are also important for stem cell behavior in invertebrates. In *Drosophila melanogaster*, reducing 3-*O* sulfation causes Notch-associated neurogenic phenotypes and lower levels of Notch protein (Kamimura et al., 2004). However, the mechanistic function of GAG chain sulfation and Notch signaling was not characterized. The fly model also showed that the glypican proteoglycans *dally* and *dally-like* are required for germline stem cell niche maintenance in females and males, respectively (Hayashi et al., 2009). These studies in combination with our work suggest that distinct embryonic and adult stem cell decisions are controlled by proteoglycan regulation of Notch expression and processing. However, the specific regulatory modality used to control Notch may depend on evolutionary origins and tissue context.

The importance of Ca^2+^ in controlling *C. elegans* germline development has previously been shown by studies of store-operated Ca^2+^ entry (SOCE) regulators - STIM-1 (stromal interaction molecule 1) and ORAI-1 (Ca^2+^ release-activated Ca^2+^ channel protein 1). Somatic-specific RNAi knockdown of either *stim-1* or *orai-1* from early larval development causes sterility in adults, suggesting a non-cell autonomous affect on the germline development (Lorin-Nebel et al., 2007; Yan et al., 2006). Our data also show that reduced germ cell proliferation caused by loss of SDN-1 is suppressed by removal of the TRP-2 Ca^2+^ channel, supporting a role for Ca^2+^ regulation of germline function. The important somatic sheath function of innexin gap junction proteins INX-8 and INX-9 in fertility suggest a possible route for communication of Ca^2+^ signals to the germline (Starich et al., 2014). In support of this, a recent study revealed that the INX-8/9- expressing sheath cell pair 1 (Sh1) is closely associated with the majority of distal germ cells and promotes their proliferation (Gordon et al., 2020). It will be interesting to investigate in subsequent studies precisely how Ca^2+^ levels are communicated from the soma to germline.

It has become clear that AP-2 transcription factors play highly conserved functions in regulating germ cell biology. Mammalian AP2γ, encoded by the Tfap2C gene, functions with additional transcription factors (BLIMP1, PRDM14 and SOX17) to promote germ cell and pluripotency genes and to inhibit somatic fates (Nakaki et al., 2013; Pastor et al., 2018; Weber et al., 2010). Further, a recent study in the cnidarian *Hydractinia symbiolongicarpus* showed that *Tfap2*, an AP2γ homolog, is essential for germ cell and gonad development (DuBuc et al., 2020). Our data show that APTF-2, a *C. elegans* homolog of AP2γ, promotes mitotic germ cell fate through direct, and Ca^2+^-dependent, regulation of the GLP-1 Notch receptor. ChIP-sequencing analysis in mammalian cells suggest that a direct regulatory relationship between AP-2 transcription factors and Notch is highly conserved. AP-2 transcription factors interact with the promoter of both Notch 1 and 3 in P19 embryonal carcinoma cells and with the Notch 2 promoter in MCF-7 breast cancer cells (Magnusdottir et al., 2013; Woodfield et al., 2010). Therefore, further study is warranted to investigate the importance of AP-2 transcription factor regulation of Notch expression in disease processes, and how proteoglycans potentially modulate Notch transcription. To conclude, our study reveals a mechanism where SDN-1 acts as a conduit for somatic-germline communication to optimize germ cell behavior and progeny generation. Future work should focus on understanding how syndecans utilise their highly complex sugar chain antennae to interpret extracellular signals to optimize germ cell fates.

## ACKNOWLEDGEMENTS

We thank Sarah Crittenden, Hannes Bulow, Alicia Melendez and John Couchman and members of the Pocock Laboratory and for advice and comments on the manuscript. Some strains were provided by the *Caenorhabditis* Genetics Center (University of Minnesota), which is funded by NIH Office of Research Infrastructure Programs (P40 OD010440). We extend our thanks Guangshou Ou and Zhiwen Zu for their expertise and advice in genome-editing to generate CRISPR/Cas9 tagged APTF-2::GFP. We also appreciate Judith Kimble and Sarah Crittenden for making the GLP-1::V5 CRISPR/Cas9 strain available prior to publication and for the kind gift of *sdn-1* Minimos rescue constructs by Hannes Bulow and Dayse S. da Cunha.

## Data and materials availability

All data is available in the main text or the supplementary materials.

## AUTHOR CONTRIBUTIONS

Conceptualization, S.G. and R.P.; Methodology, S.G. and R.P.; Investigation, S.G., A.A., A.E., L.N. and R.P. Writing - Original Draft, R.P.; Writing - Review & Editing, S.G., A.A., L.N. and R.P.; Funding Acquisition, S.G., R.P.; Resources, S.G. and R.P; Supervision, S.G. and R.P.

## STAR Methods

### CONTACT FOR REAGENT AND RESOURCE SHARING

Strains used in this study will be deposited at the *Caenorhabditis* Genetics Center (CGC) and will be available upon request. Further information and requests for resources and reagents should be directed to and will be fulfilled by the Lead Contacts Roger Pocock and Sandeep Gopal.

### EXPERIMENTAL MODELS AND SUBJECT DETAILS

#### Caenorhabditis elegans

*C. elegans* strains were cultured on Nematode Growth Medium (NGM) plates and fed with OP50 *Escherichia coli* bacteria at 20°C, unless otherwise stated. All strains used in this study are listed in the Key Resources Table. Experiments were performed in triplicates and the number of animals analyzed is annotated in each Figure legend.

#### Mammalian cells

For Electrophoretic Mobility Shift Assays, HEK293T cells were cultured in minimal essential media with 10% FCS and 2.5mM glutamine.

##### Generation of transgenic strains

Transgenic lines were generated by injecting DNA constructs into young adult hermaphrodites as complex arrays. *sdn-1* rescue lines were generated using miniMos-mediated genomic insertion as previously described (Frokjaer-Jensen et al., 2014). Briefly, *sdn-1(zh20)* animals were injected with a plasmid mix containing *rab-3p::mCherry::unc-54UTR, myo-2p::mCherry::unc-54UTR, myo-3p::mCherry::unc-54UTR, eft-3p::mos1transposase::tbb-2UTR, hsp16.41p::peel-1::tbb-2UTR* and *sdn-1::gfp::tbb-2UTR* driven by specific promoters (kind gifts of Hannes Bulow). The promoters used to drive *sdn-1* expression were *lag-2p* (DTC), *mex-5p* (germline) and *lim-7p* (gonadal sheath). Injected worms were screened for expression of fluorescent co-injection markers and positive worms were treated with 500μl G418 antibiotic (25μg/μl). 14 days after antibiotic treatment, worms were heat shocked at 34°C for 3hr. Surviving worms were picked individually to new plates to establish independent lines.

##### Generation of strains using CRISPR-Cas9

###### Endogenous tagging of APTF-2 with GFP using CRISPR-Cas9

A C-terminal GFP knock-in strain for endogenous *aptf-2* was generated using CRISPR/Cas9-triggered homologous recombination (Dickinson et al., 2013). The Cas9-sgRNA construct of *aptf-2* was obtained using PCR to insert the target sequence: (GCCAACTGAATCAAAGCCAGAGG) into pDD162 (Addgene #47549). For the GFP tag knock-in construct, homology recombination templates were constructed with the following steps: 1) cloning a 2kb genomic region of *aptf-2* centered on the knock-in site into the pPD95.75 vector, 2) insertion of DNA encoding GFP and a linker of six amino acids (AGACCCAAGCTTGGTACC) between the last amino acid codon of *aptf-2* and the stop codon, 3) the PAM site within the recombination template was mutated to prevent Cas9 cleavage. The following mix was then injected into wild-type animals: 10 ng/μl homologous knock-in repair template, 10 ng/μl Cas9-sgRNA plasmid, *myo-2::mCherry* plasmid (4 ng/μl). Individual F1 progeny of injected wild-type worms were picked to individual plates and F2 progeny screened for GFP knock-in by PCR. After confirmation of insertion by Sanger sequencing, the GFP knock-in was outcrossed three times prior to analysis. APTF-2::GFP was imaged in 1-day old adult germlines and 2-cell stage embryos using a Zeiss Axiocam 40x objective.

##### *glp-1::V5* promoter mutations

Repair oligos and injections were performed essentially as described by injecting into *glp-1::V5(q1000)* animals (Dokshin et al., 2018). To generate mutations at the APTF-2 target site in the *glp-1* promoter we injected the following mix: Cas9 protein (5μg), tracrRNA (0.4μg/μl), crRNA (aatgggcggagtatgccacc) (0.4μg/μl), ssDNA oligo repair donor (1μg/μl), *myo-2::mCherry* plasmid (4ng/μl) and brought to a 20μl volume with nuclease-free water. Mutation/deletion of the APTF-2 binding site was screened by PCR followed by restriction digest with *NciI*. Mutations were verified by Sanger sequencing and outcrossed three times prior to analysis.

##### Staining and germline analysis

Semi-automated germline analysis was performed as previously reported (Gopal et al., 2017). L4 hermaphrodites were picked to OP50 plates and incubated for 16hr at 20°C to reach the young adult stage. Germlines were extruded from sedated worms and fixed on a poly-L-lysine coated slides using ice cold methanol for 1min and then in 3.7% paraformaldehyde (PFA) for 25min. Fixed germlines were washed three times in phosphate buffered saline (PBS, pH 7.4) and blocked using 30% normal goat serum before incubating with primary antibodies overnight at 4°C. After incubation, germlines were washed three times with PBS containing 1% Tween-20 (PBST) and incubated with fluorophore-conjugated secondary antibodies and 4′,6-diamidino-2-phenylindole (DAPI) for 1hr at 25°C. After staining, germlines were washed three times with PBST. Slides were mounted by applying a drop of Fluroshield mounting media (Sigma) on the germlines followed by a coverslip. Stained germlines were analyzed using either the Zeiss Axiocam 40x objective (GLP-1-V5 staining and smFISH) or the Leica SP5 63x objective (scoring germ cell number). GLP-1-V5 expression and smFISH fluorescence was quantified using FIJI according the formula, Corrected Total Fluorescence (CTF) = Integrated Density – (Area of the mitotic region x Mean fluorescence of background readings). Germ cell number and 3D modelling were performed using Imaris 9.5. Briefly, stained germlines were imaged with confocal microscopy at 63x magnification and additional 1.7x zoom. The diameter of DAPI stained nuclei in the mitotic region was defined as 2.5μm and a three-dimensional model was created to enable counting of nuclei. To quantify the number of cells in active mitosis, extruded germlines were stained with an anti-pH3 antibody and pH3-positive nuclei counted.

##### Transfection of HEK293T cells

HEK293T cells were seeded on a 3-cm dish one day prior to transfection. Cells were transfected at 80% confluency using Lipofectamine 2000 (Thermofisher) Briefly, 2μg pcDNA3.2-*aptf2::V5* was diluted in 200μl Opti-MEM (Life Technologies) and 8μl Lipofectamine was diluted in 200μl Opti-MEM. Both solutions were incubated separately at 25°C for 5min before being combined for a further incubation at 25°C for 20min. The mixture was added to the dish containing cells in 1ml of media and incubated for 24hr at 37°C prior to harvesting for nuclear preparation and Electrophoretic Mobility Shift Assay.

##### Biotin Labelling and DNA duplex formation

60 bp oligonucleotides encompassing part of the *glp-1* promoter (wild-type or with the TFAP2 consensus sequence deleted - AP-2 del) (Key Resources Table) were labelled using a 3’ end DNA biotin labelling kit (Thermo Scientific). First, 1μM oligonucleotides were incubated with 0.12U/μl Terminal Deoxynucleotidyl Transferase, 1x Terminal Deoxynucleotidyl Transferase buffer (500mM cacodylic acid, 10mM CoCl_2_, 1mM DTT, pH 7.2) and 0.5μM Biotin-11-UTP at 37°C for 30min. The reaction was stopped by adding 2.5μl of 0.2M EDTA and labelled oligonucleotides extracted from the reaction mix by adding 50μl chloroform:isoamyl alcohol. Biotin labelled complementary oligonucleotides were incubated together at 90°C for 2min. The temperature was gradually reduced to 70°C and incubated for 30min before further reducing the temperature to 10°C. This resulted in the formation DNA duplexes with biotin labels at the 3’ ends.

##### Electrophoretic Mobility Shift Assay (EMSA)

HEK293T cells (see above) were washed with PBS and nuclear fractions extracted using NE-PER Nuclear and Cytoplasmic Extraction Reagents (Thermo Scientific). Briefly, 50μl of packed volume of cells was lysed with 500μl of cytoplasmic extraction reagent I and 27.5μl of cytoplasmic extraction reagent II to remove the cytoplasmic fraction. The remaining cell pellets were treated with 500μl of nuclear extraction reagents for 40min and the solution was centrifuged at 16000rpm for 5min. The supernatant containing the nuclear fraction was collected for EMSA. EMSA was performed according to the instructions from the LightShift® Chemiluminescent EMSA Kit (Thermo Scientific). 8μl of the nuclear fraction was incubated with 15nM of biotin labeled DNA duplex, 2.5% glycerol, 5mM MgCl_2_, 0.05% NP-40 and 50 ng/μl Poly(deoxyinosinic-deoxycytidylic) acid for 25min at 25°C. Three sets of reactions were prepared: 1) APTF-2::V5 nuclear fraction and biotin labelled wild-type *glp-1* DNA duplex, 2) APTF-2::V5 nuclear fraction and biotin labelled *glp-1* AP-2 del DNA duplex and 3) biotin labelled wild-type *glp-1* DNA duplex alone. Another set of competition reaction mixes were prepared by adding incremental amounts of unlabelled wild-type *glp-1* DNA duplex in addition to biotin labelled wild-type *glp-1* DNA duplex. For all reactions, an equal amount of APTF-2 nuclear fraction was used. After a 30min incubation, 5μl of 5x loading buffer was added to the reaction mix. Samples were then run on a native polyacrylamide gel in 1x Tris-Borate-EDTA buffer. Gels were blotted onto nitrocellulose membranes. The membranes containing DNA were crosslinked using a UV transilluminator for 15min, followed by incubation with blocking buffer provided in the EMSA kit for 15min at 25°C. Then, the membrane was incubated with blocking buffer containing stabilized Streptavidin-Horseradish Peroxidase Conjugate (1:300) for 15min followed by five washes with wash buffer. The membrane was then incubated in substrate equilibration buffer for 15min and treated with chemiluminescent substrate for 5min. The membrane was exposed using Biorad ChemiDoc XRS+ and images were taken using a CCD camera and Image lab 6.0.1 software. To examine the effect of Ca^2+^ on APTF-2 binding to the *glp-1* promoter, the EMSA was performed with the reaction mix containing 100nM of CaCl_2_. Controls were performed by using a reaction mix containing Epstein-Barr Nuclear Antigen Extract (1U) and 60 bp biotin end-labeled duplex (20fmol) containing the following binding site 5’-TAGCATATGCTA-3’.

##### Single-Molecule Fluorescence in Situ Hybridization (smFISH)

smFISH was performed using Stellaris RNAi FISH reagents (Biosearch Technologies). Twenty young adult hermaphrodites were grown in OP50 plates to lay embryos. After 4hr, worms were removed from the plates and embryos were washed three times with M9 buffer to remove any bacteria before being fixed using 3.7% PFA in PBS for 45min at 25°C. Embryos were then permeabilized using 70% ethanol at 4°C for 16hr followed by an incubation with Buffer A at 25°C for 5min. The *glp-1* FISH probes were prepared in 100μl of the Hybridization Buffer and worms were incubated in the probe solution in the dark at 37°C for 4hr. 500μl Buffer A was added to the embryos and incubated at 37°C for 30minutes followed by washing with Buffer A. Samples were then incubated with DAPI stain solution (Wash Buffer A consisting of 5 ng/ml DAPI). After a 30-min incubation in the dark at 37°C, the DAPI solution was removed and embryos were washed with Buffer B. Embryos were transferred onto a slide containing mounting media and a coverslip was placed on top. Images were captured using the Zeiss Axiocam 40x objective and fluorescence intensity was measured using FIJI.

##### RNA interference

HT115(DE3) *E. coli* bacteria expressing RNAi plasmids for specific genes or empty vector (L4440) were grown in Luria Broth (LB) + Ampicillin (50μg/ml) at 37°C for 16hr. Saturated cultures of RNAi bacteria were plated on RNAi plates and allowed to dry for 24hr. L4 hermaphrodites were placed on the RNAi plates and incubated for 16hr at 20°C before proceeding with germline analysis.

##### Brood size analysis

10 L4 hermaphrodites were picked onto individual NGM plates seeded with OP50 bacteria. Worms were allowed to lay eggs for 24hr and then the mothers were individually moved to new plates. After a further 24hr embryos were analyzed for hatching. This process was repeated for six days. The number of larvae and embryos were counted each day and summed as the total brood size.

##### Western blotting

Transgenic worms expressing GFP-tagged APTF-2 were grown on NGM plates coated with OP50 bacteria. A packed volume of 200μl of mixed stage worms and embryos were collected by washing with M9 buffer. Worms were washed three times with M9 buffer, pelleted by centrifugation and 250μl of lysis buffer (50mM Tris pH 7.4, 150mM NaCl, 2% triton X-100, 0.1% SDS, 1x protease inhibitor cocktail) was added. The samples were sonicated using Bioruptor^®^ (Diagenode) at high amplitude (4°C; 30 sec ON and 20 sec OFF) for 10-15 cycles. Worms were then disrupted using a mortar and pestle for 5min before centrifuging at 3000rpm for 5min. The supernatant was collected and LDS sample buffer (1x final concentration) added before boiling at 95°C for 10min. Samples were cooled to room temperature and run on a 10% polyacrylamide gel. The gel was blotted to a PVDF membrane using the iBlot semi-dry blot system (Thermoscientific). After blocking with 5% BSA in PBS for 1hr, the PVDF membranes were incubated with anti-GFP antibody (Roche) and α-tubulin (Developmental Studies Hybridoma Bank) in 1% BSA in PBS-tween (0.1%) for 16hr at 4°C. The membrane was then washed three times with PBS-tween and incubated with horseradish peroxidase-conjugated secondary antibodies in 1% BSA in PBS-tween for 1hr at 22°C. The membrane was washed five times with PBS-tween and incubated with ECL reagents (Thermoscientific) for 5min at 22°C. The membrane was exposed using Biorad ChemiDoc XRS+ and images taken using a CCD camera and Image lab 6.0.1 software.

##### cDNA preparation and quantitative PCR

200 L4 stage hermaphrodites were grown on NGM plates for 16hr and the adult germlines were extruded. Total RNA from extruded germlines were isolated using the Qiagen RNAeasy kit. First, germlines were placed in 350μl of lysis buffer and snap-frozen in liquid nitrogen before being thawed at 37°C. Freeze/thaw cycles were repeated seven times and an equal volume of 70% ethanol was added. The samples were transferred to RNAeasy spin columns and centrifuged at 13000rpm for 15 sec to remove flow through. In order to eliminate DNA contamination, on-column DNA digestion was performed by adding DNase in RDD buffer. Samples were then washed twice with wash buffers followed by 80% ethanol before being eluted in 14μl of RNAase free water. cDNA was synthesized from the total RNA using the ImProm-II™ Reverse Transcription System. Briefly, 500 ng of RNA was mixed with 5μl of oligo-dT (0.05μg/μl) at 70°C for 5min and then 15μl of master mix (1x ImProm II buffer, 0.05mM dNTP, 20U RNasin Ribonuclease inhibitor, 2.5mM MgCl_2_ and 1U ImProm II) was added. cDNA was synthesized in a thermocycler (5min at 25°C, 1hr at 42°C, 15min at 70°C). To perform qPCR, 1μl of cDNA was mixed with 5μl of SyBR master mix (Roche), 1μl of forward primer (10μM) and 1μl reverse primer (10μM) on 96 well plates. Reactions were run on a LightCycler 480 II real time PCR machine for 35 cycles. Two housekeeping genes, *cdc-42* and *pmp-3,* were used as reference gene controls. The Ct values obtained were used to calculate the mRNA levels and values were normalized against values obtained from the reference genes. Graphs were plotted as a fold change of *glp-1* mRNA in *sdn-1* mutants with respect to the wild-type worms.

##### Site-directed mutagenesis for SDN-1 construct (S-A mutations)

Three heparan sulfate chains attach to the SDN-1 core protein at ^71^Ser, ^86^Ser and ^214^Ser. In order to prevent heparan sulfate chain attachment to the core protein, these serine residues were substituted with alanine using the Q5^®^ Site-Directed Mutagenesis Kit. ^71^Ser and ^86^Ser were substituted in a single reaction using the oligos oSA1F and oSA1R (Key Resources Table). The plasmid containing *lim-7p::sdn-1::gfp::tbb-2UTR* was used as the template for the first two substitutions. The resulting plasmid after the first substitution was used for mutating ^214^Ser, using the primers oSA2F and oSA2R (Key Resource Table). For both mutations, a reaction mix containing 25ng of template DNA, 1x Q5 Hot Start High-Fidelity 2X Master Mix, 0.5μM forward primer and 0.5μM reverse primer was prepared and run on a thermocycler. After initial denaturation at 98°C for 30 sec, 25 cycles of 98°C for 10 sec, 62°C for 20 sec and 72°C for 5min were run. The reaction ended with a final extension for 2min at 72°C. The new plasmid was then transformed, purified and the substitutions confirmed by Sanger sequencing.

##### APTF-2-V5 expression mouse embryonic fibroblasts

Wild-type and syndecan-4 knockout mouse embryonic fibroblasts (MEFs) were transfected with APTF-2::V5 as explained above. Cells were incubated for 48hr after transfection at 37°C before fixing with 4% PFA in PBS. Cells were then incubated with anti-V5 antibody at 4°C overnight before incubating with fluorophore-tagged secondary antibody, DAPI and phalloidin. Cells were imaged using the Zeiss Axiocam 40x objective.

### QUANTIFICATION AND STATISTICAL ANALYSIS

All experiments were performed in three independent replicates and the experimenter was blinded to genotype. Statistical analysis was performed in GraphPad Prism 7 using one-way analysis of variance (ANOVA) for comparison followed by Dunnett’s Multiple Comparison Test where applicable. Welch’s t-test was performed if the comparison was for two conditions. Values are expressed as mean ± s.e. Differences with a *P* value <0.05 were considered significant.

## SUPPLEMENTARY FIGURES

**Figure S1. *sdn-1* loss-of-function germline analysis**

(A) Brood size of wild-type, *sdn-1(zh20)* and *sdn-1(ok449)* hermaphrodites. Data are expressed as mean ± s.e.m. and statistical significance was assessed by ordinary one-way ANOVA. n>11. ****p<0.0001, *** p<0.001.

(B) PZ length of wild-type, *sdn-1(zh20)* and *sdn-1(ok449)* adult hermaphrodites. Data are expressed as mean ± s.e.m. and statistical significance was assessed by ordinary one-way ANOVA. n>40. ****p<0.0001.

(C) 3D germline quantification of PZ nuclei in adult hermaphrodite (+/− *sdn-1* RNAi). Data are expressed as mean ± s.e.m. and statistical significance was assessed by unpaired t-test. n>15. ****p<0.0001.

**Figure S2. *sdn-1* mutant rescue experiments**

(A) Quantification of M-phase cells in distal germlines of wild-type, *sdn-1(zh20)* and *sdn-1(zh20); lim-7p::sdn-1 cDNA* adult hermaphrodites. The *lim-7p::sdn-1 cDNA* rescue line corresponds to the same line in Figure 1F. Data are expressed as mean ± s.e.m. and statistical significance was assessed by ordinary one-way ANOVA. n=40. ***p<0.001.

(B) Schematic of wild-type SDN-1 and GAG-deficient SDN-1. Attachment sites (S71, S86 and S214) are shown with GAG chains as red lines in wild-type. Alanine mutations that prevent GAG attachment are shown in the GAG-deficient SDN-1. Yellow - signal sequence; White - ectodomain; Green - transmembrane domain; Grey - cytoplasmic domain.

(C) Quantification of PZ length in distal germlines of adult hermaphrodites in wild-type, *sdn-1(zh20)* and *sdn-1(zh20); lim-7p* rescue lines expressing wild-type SDN-1 or GAG-chain deficient SDN-1(AAA). Independent rescue lines represented by # (#1 line for wild type SDN-1 corresponds to the same line in Figure 1F). Data are expressed as mean ± s.e.m. and statistical significance was assessed by ordinary one-way ANOVA. n=40. ****p<0.0001, n.s. not significant.

(D) Quantification of M-phase cells in distal germlines of adult hermaphrodites in wild-type, *sdn-1(zh20)* and *sdn-1(zh20); lim-7p* rescue lines expressing wild-type SDN-1 or GAG-chain deficient SDN-1(AAA). Independent rescue lines represented by # (#1 line for wild type SDN-1 corresponds to the same line in Figure 1F). Data are expressed as mean ± s.e.m. and statistical significance was assessed by ordinary one-way ANOVA. n=40. ****p<0.0001, n.s. not significant.

**Figure S3. *glp-1* epistasis analysis**

(A) Quantification of PZ nuclei in wild-type, *sdn-1(zh20)*, *glp-1(e2141)* and *sdn-1(zh20)*; *glp-1(e2141)* adult hermaphrodites. Data are expressed as mean ± s.e.m. and statistical significance was assessed by ordinary one-way ANOVA. n>22. ****p<0.0001, n.s. = not significant.

(B) Quantification of M-phase cells in distal germlines in wild-type, *sdn-1(zh20)*, *glp-1(e2141)* and *sdn-1(zh20)*; *glp-1(e2141)* adult hermaphrodites. Data are expressed as mean ± s.e.m. and statistical significance was assessed by ordinary one-way ANOVA. n>23. ****p<0.0001, n.s. not significant.

**Figure S4. Anaysis of CRISPR-Cas9 generated GLP-1::V5 strain**

(A) Quantification of PZ nuclei of wild-type and CRISPR-Cas9-tagged *glp-1::v5* adult hermaphrodites. Data are expressed as mean ± s.e.m. and statistical significance was assessed by unpaired t-test. n>20. n.s. = not significant.

(B) Images of GLP-1::V5 in adult germlines and embryos. Scale bars = 25μm.

**Figure S5. *glp-1* promoter analysis**

(A-B) *glp-1* smFISH images (A) and quantification (B) of *glp-1* transcripts (exon probes) in wild-type and *glp-1prom*Δ two-cell embryos. *glp-1* transcripts = magenta. Inset images = DAPI staining showing chromosomes of two-cell embryos. Data are expressed as mean ± s.e.m. and statistical significance was assessed by unpaired t-test. n>20. *p<0.05. Scale bar = 20μm.

(C) Quantification of GLP-1::V5 protein in wild-type*, glp-1prom(aa)* and *glp-1prom*Δ adult hermaphrodite germlines. Data are expressed as mean ± s.e.m. and statistical significance was assessed by ordinary one-way ANOVA. n>40. ****p<0.0001, ***p<0.001.

(E) Quantification of PZ nuclei in wild-type*, glp-1prom(aa)* and *glp-1prom*Δ adult hermaphrodites. Data are expressed as mean ± s.e.m. and statistical significance was assessed by ordinary one-way ANOVA. n>40. ****p<0.0001.

**Figure S6. PZ nuclei analysis of *aptf-1* and *aptf-2* mutants**

Quantification of PZ nuclei in wild-type, *aptf-1(gk794)* and *aptf-2(qm27).* n>15. ****p<0.0001, n.s. = not significant.

**Figure S7. *glp-1* EMSA analysis**

(A-B) EMSA competition experiments. APTF-2::V5 protein produced in HEK293T cells and the TFAP2 consensus motif in the *glp-*1 promoter (A). DNA bound and unbound with APTF-2::V5 is marked. Biotin-labelled wild-type *glp-1* promoter is competed away with increasing amount of unlabeled *glp-1* promoter. Quantification of shifted *glp-1* promoter with increasing amount of unlabeled *glp-1* promoter (B).

**Figure S8. APTF-2::GFP expression analysis**

(A) *aptf-2* genomic locus (upper image) showing 3’-end tagging of the gene (crRNA used for Cas9 cleavage in red). Schematic of APTF-2 protein (lower image) showing approximate molecular weight of the fusion protein following CRISPR/Cas9 insertion of GFP.

(B) Western blot of endogenously tagged APTF-2::GFP from mixed-stage worms. APTF-2::GFP protein is the correct size (APTF-2 = ~40kDa, GFP = 27kDa, combined = ~67kDa). Top blot - anti-GFP antibody; bottom blot - anti-tubulin loading control.

(C-D) Endogenous APTF-2::GFP expression in the adult hermaphrodite germline. DIC image of a young adult hermaprhodite (C), fluorescence image of the same worm (D) and zoomed image of the boxed region (D’) showing expression in the germline (white arrowheads) and oocytes (yellow arrowheads). Scale bar = 50μm.

